# Effects of combined use of alcohol and delta-9-tetrahydrocannibinol on working memory in Long Evans rats

**DOI:** 10.1101/2023.02.02.526698

**Authors:** Lauren K. Carrica, Chan Young Choi, Francis A. Walter, Brynn L. Noonan, Linyuan Shi, Clare T. Johnson, Heather B. Bradshaw, Nu-Chu Liang, Joshua M. Gulley

## Abstract

The increase in social acceptance and legalization of cannabis over the last several years is likely to increase the prevalence of its co-use with alcohol. In spite of this, the potential for effects unique to co-use of these drugs, especially in moderate doses, has been studied relatively infrequently. We addressed this in the current study using a laboratory rat model of voluntary drug intake. Periadolescent male and female Long-Evans rats were allowed to orally self-administer ethanol, Δ^9^-tetrahydrocannibinol (THC), both drugs, or their vehicle controls from postnatal day (P) 30 to P47. They were subsequently trained and tested on an instrumental behavior task that assesses attention, working memory and behavioral flexibility. Similar to previous work, consumption of THC reduced both ethanol and saccharin intake in both sexes. Blood samples taken 14h following the final self-administration session revealed that females had higher levels of the THC metabolite THC-COOH. There were modest effects of THC on our delayed matching to position (DMTP) task, with females exhibiting reduced performance compared to their control group or male, drug using counterparts. However, there were no significant effects of co-use of ethanol or THC on DMTP performance, and drug effects were also not apparent in the reversal learning phase of the task when non-matching to position was required as the correct response. These findings are consistent with other published studies in rodent models showing that use of these drugs in low to moderate doses does not significantly impact memory or behavioral flexibility following a protracted abstinence period.

---

Alcohol and cannabis are two of the substances most used by adolescents, with about 41% of 18-year-olds estimated to have consumed alcohol within their lifetime and about 30% having used cannabis (Johnston et al., 2019). Deficits in executive function, attention span, working memory, and other aspects of cognition have previously been identified in those who initiated moderate to heavy use of alcohol (Squeglia et al., 2009; Mota et al., 2013; Mahedy et al., 2018) or cannabis (Harvey et al., 2007; Medina et al., 2007; Hanson et al., 2010; Hooper et al., 2014) during adolescence. Although it is often the case that these drugs are taken simultaneously, especially in late adolescents/young adults (McCabe et al., 2021), the consequences of this behavior have been studied less frequently than their independent use (Subbaraman and Kerr 2015; Yurasek et al., 2017). When simultaneous use is studied, it is often in terms of immediate effects on intoxication and drug-taking patterns. For example, in young adults who co-use alcohol and cannabis, there is a tendency to drink more alcohol compared to when cannabis is not used, as well as an increased incidence of negative consequences of drug use (Jackson et al., 2020; Linden-Carmichael et al., 2020; Linden Carmichael et al., 2019).

Studies in laboratory rodents have largely focused on the effects of one of these drugs in isolation. As in the human literature, cognitive deficits have been demonstrated in rat and mouse models of adolescent exposure to ethanol or Δ^9^-tetrahydrocannibinol (THC; for reviews of this literature, see Spear, 2018; Bara et al., 2021). In some studies, these effects are more pronounced in adolescents compared to adults (Hunt and Barnet, 2016;), whereas others find no significant effects or even improvement of cognitive function following alcohol or cannabinoid exposure (Bruijnzeel et al., 2019; Boutros et al., 2017; Slawecki, 2006; Broadwater and Spear, 2013). In the limited number of cases where co-exposure to these drugs has been studied, results have been mixed. Using forced exposure to vaporized THC combined with voluntary alcohol drinking, Hamidullah and colleagues (2021) found modest deficits in recognition memory and instrumental learning for adult rats that were alcohol- or THC-exposed from postnatal days (P) 28-42. These effects were similar or absent in rats concurrently exposed to both drugs. In rats exposed to vaporized THC from P45-50 and vaporized ethanol from P50-64, Smiley et al. (2021) found evidence for increased freezing responses in a conditioned fear paradigm in rats exposed to THC, ethanol, or both drugs. These behavioral effects were not different between groups of drug-exposed rats, but rats exposed to both drugs uniquely expressed an enhanced calcium signaling response in the prelimbic cortex to footshocks delivered in the last half of the fear conditioning test session. In studies from our group that investigated the effects of voluntary THC and ethanol exposure from P30-45 on spatial memory, behavioral flexibility, and abstinence-related anhedonia, we found no evidence for long-term effects of peri-adolescent drug exposure on behavior (Nelson et al., 2019a). These previous studies of co-exposure effects utilized only male rats despite known sex differences in the effects of alcohol alone or THC alone in humans (Thomasson, 1995; Cooper and Craft, 2018) and laboratory rodents (Knott et al., 2015; Rubino et al., 2008). Importantly, there is evidence that the incidence of alcohol and cannabis co-use is significantly higher in men compared to women (Subbaramen and Kerr, 2015; Patrick et al., 2019; Briere et al., 2011). Furthermore, female adolescents who co-use alcohol and cannabis have been reported to have higher levels of impairment in working memory than their male counterparts in both cannabis-only and combined alcohol and cannabis groups (Noorbakhsh et al., 2020).

In the current study, we utilized a rat model of peri-adolescent alcohol and THC co-use in both male and female rats to investigate the potential effects of combined consumption on working memory performance in adulthood. Through the use of a model of THC consumption previously development by our group (Nelson et al., 2019a,b), we aimed to model a method of human drug use through THC-laced food products (i.e., “edibles”). Recent reports have suggested that edible use may be on the rise with increasing legalization of medicinal and recreational use cannabis, especially among adolescents (Patrick et al., 2020). Rats were allowed to self-administer either 10% ethanol with 0.1% saccharin, THC, or both ethanol and THC from P30 to P47. When they reached young adulthood (P82), rats were tested in a delayed matching to position task (DMTP) that assess working memory. Based on the findings in human literature (Noorbakhsh et al., 2020; Linden-Carmichael et al., 2020 we hypothesized that combined alcohol and THC use during adolescence would impair working memory in adulthood to a greater extent than exposure to either drug alone. Due to the relatively moderate level of exposure to one or both drugs, we anticipated to see these differences in only the more challenging aspects of the task, e.g. when the delays were long and upon rule reversal in the DNMTP Task. This a priori hypothesis, as well as the methods and data analysis plan for this study, were pre-registered at the Open Science Framework prior to the start of data collection (https://osf.io/tjxvd/).

## Methods

### Subjects

Long-Evans rats (72 male, 60 female) were obtained from Envigo (Indianapolis, IN) and delivered to our colony at approximately postnatal day (P) 24 (±2 days). Rats were housed in same-sex pairs in a temperature-controlled room on a 12:12 light/dark cycle (lights on at 1000 h) with food and water available ad libitum until the beginning of behavioral training as described below. Volitional exposure to drugs or vehicle took place during portions of the dark and light phases of the cycle; operant behavior sessions all took place during the light phase. Rats were pair-housed with same-sex rats and were weighed daily starting on the day following arrival. Though we had planned at the outset of the study for three cohorts, a fourth was added due to attrition (criteria for study removal described below). Between 11 -14 rats from each sex that were assigned to each of the four treatment groups met drug-taking and behavioral performance criteria to be included in this report (51 male, 53 female). All experimental procedures were performed in accordance with the Guide for the Care and Use of Laboratory Animals (National Research Council, 2011) and were approved by the Institutional Animal Care and Use Committee (IACUC) at the University of Illinois, Urbana-Champaign.

### Drugs and exposure procedures

We utilized a similar procedure from our previous studies investigating volitional exposure to ethanol and THC (Nelson et al., 2018). THC (200 mg/mL dissolved in 95% ethanol; Research Triangle Institute; distributed by The National Institute on Drug Abuse) was diluted in 95% ethanol (Decon Laboratories) and then directly applied via pipette to halves of cookies (Pepperidge Farm Fudge Brownie Goldfish Grahams). The 95% ethanol vehicle was applied to a separate batch of cookie halves for rats assigned to receive a control (vehicle) cookie. THC doses were given following the schedule outlined below. Ethanol, saccharin (saccharin sodium salt hydrate; Sigma-Aldrich), and tap water were used to prepare 0.1% saccharin solutions and 0.1% saccharin solutions in 10% ethanol. The caloric free, sweet saccharin solution was used to increase drinking during the scheduled ethanol access period without interfering with caloric satiety.

The treatment procedure is summarized in Table 1. Rats were assigned randomly to one of four treatment groups: control (CTL), which were given access to 0.1% saccharin solutions (SAC) and vehicle-laced cookies; ethanol only (EtOH), which were given access to 10% EtOH/SAC solutions and vehicle-laced cookies; THC only (THC), which were given access to SAC solutions and THC-laced cookies; and the combined exposure group (COM), which were given access to 10% EtOH/SAC solutions and vehicle-laced cookies. Starting on P26 and continuing until the end of vehicle or drug exposure (P47), cagemates were physically separated from each other by insertion of a clear acrylic divider into their homecage. This divider, which contained small holes to allow for potential olfactory and tactile interaction between cagemates, allowed for individual presentation of fluid or cookies and measurement of individual intake while maintaining many aspects of social housing. These dividers were removed following the last day of drug exposure; they were re-inserted in < 10% of cages when instances of aggressive behavior between cagemates were noted.

**Table 1.** Schedule for presentation of fluid [0.1% saccharin (SAC) or 10% ethanol + SAC (EtOH)] and cookies [vehicle (95% ethanol) or THC laced]. Daily THC doses are increased every 4 – 6 days and delivered on two occasions as indicated in the rows of Cookies 1 and 2, which together targeted daily doses of 3, 6, 8, and 10 mg/kg.

| Time/Age (P) | 28 – 29 | 30 – 31 | 32 – 35 | 36 – 39 | 40 – 43 | 44 – 47 |
| --- | --- | --- | --- | --- | --- | --- |
| 07:30 – 10:00 h<br>(dark cycle) | SAC | SAC | EtOH or<br>SAC | EtOH or<br>SAC | EtOH or<br>SAC | EtOH or<br>SAC |
| Cookie 1<br>(~ 10:00 h) | vehicle | 1.5 THC<br>or vehicle | 1.5 THC<br>or vehicle | 3.0 THC<br>or vehicle | 3.0 THC<br>or vehicle | 5.0 THC<br>or vehicle |
| Cookie 2<br>(~ 19:30 h) | vehicle | 1.5 THC<br>or vehicle | 1.5 THC<br>or vehicle | 3.0 THC<br>or vehicle | 5.0 THC<br>or vehicle | 5.0 THC<br>or vehicle |

On P28 and P29, rats were provided access to sipper tubes containing the SAC solution during the last 2.5 h of the dark cycle. At light onset, fluid intake and body weight were measured after which the first vehicle (25 μL of 95% ethanol)-laced cookie was given. About 10 h later and 2.5 h prior to the onset of the dark cycle, the second vehicle-laced cookie was given. Rats quickly established significant intake of SAC and complete consumption of cookies. After this training, rats were assigned to one of the four groups so that mean baseline fluid intake and body weight was similar between groups. THC exposure began on P30 when CTL and EtOH rats continued to receive vehicle-laced cookies throughout the 18-day exposure period, whereas the THC and COM rats received THC-laced cookies following the dosing schedule outlined in Table 1. Two days following the onset of THC exposure, fluid access for EtOH and COM rats was switched to 10% ethanol (v/v) in SAC whereas the CTL and THC rats continued to receive the SAC solution until the end of drug exposure. We previously reported that rats drinking alcohol using the same protocol used here reached an average blood alcohol level of ∼ 30 mg/dL when measured three hours after drinking (Nelson et al., 2019a). Using LC-MS/MS to measure THC and its metabolites, our pilot study indicated that blood levels of THC reached after consuming a 5 mg/kg THC-laced cookie were at least 10-fold lower than what has been associated with cannabis dependence in rats (Bruijnzeel et al., 2016). Thus, the ethanol and THC intake in this paradigm was considered moderate exposure. Ad libitum access to tap water in a standard water bottle was available to all rats during the remaining hours of the day. Similarly, ad libitum access to standard chow (3.1 kcal/g; 58% carbohydrate, 24% protein, and 18% fat; Envigo 2018 Teklad Global rodent diet, Indianapolis, IN, USA) was available to all rats.

### Plasma Levels of THC and Metabolites

Tail blood was collected approximately 14h after the final THC exposure in both THC and COM groups. Samples were quickly centrifuged for plasma extraction, and plasma was stored at -80 °C until analysis. Liquid chromatography-tandem mass spectrometry (LC-MS/MS) was used to quantify levels of THC, 11-OH-THC, and THC-COOH, as previously described (Leishman et al., 2018).

### Apparatus

Operant behavior was assessed in standard modular chambers (Coulbourn Instruments). The front wall of each chamber contained a centrally located food trough, a retractable lever on either side of the trough (i.e., the right or left lever), and a white cue light above each lever. The back wall contained a white houselight near the top of the chamber and a recessed nosepoke port containing a red LED light that was located 1.8 cm above the floor. Infrared photobeam detectors located inside the food trough and nosepoke port were used to monitor head entries. Graphic State (v4.1; Coulbourn Instruments) was used for automated chamber control as well as data collection.

### Working Memory Task

The task is a modified version from Dunnett and colleagues (Sloan et al., 2006) that we have used previously (Sherrill et al. 2013; Hankosky et al. 2017). On P54, rats’ access to food was restricted to a once daily allotment that was designed to reduce body weight over five days to 80-90% of their *ad-libitum* feeding weight. To account for the significant body weight change that typically occurs as young, free-fed Long Evans rats mature, the target weight range for each rat was adjusted daily through P70 so that body weight was maintained within 80-90% of an age- and sex-matched, free-fed “control” rat using normative growth curve data provided by the vendor.

Beginning at P59, rats were trained over successive days to respond on either the left or right lever to earn a food pellet (45 mg, BioServ F0021) on an FR1 schedule. Rats were then trained to perform a matching-to-position (MTP) task using a forward chaining approach, which involves training on each progressive step of a multi-part sequence such that rats eventually learn all components of the task. An individual trial began with a 5-sec intertrial interval (ITI), followed by extension of one of the levers (selected randomly) and the illumination of the cue light above the lever. A lever press within this 20-sec “sample phase” led to illumination of the nosepoke port on the opposite wall and the lever retracted. If the rat poked its nose into the port on the back wall within 10 sec, the “choice phase” began: the port light turned off, both levers extended, and the cue lights above the levers illuminated. If the rat responded on the lever previously presented during the sample phase within 10 sec, a food pellet was delivered into the trough (illuminated for 3 sec) and the levers retracted. A response on the non-sample lever was scored as incorrect. A failure to respond on the levers or at the backwall nosepoke at any phase was counted as an omission.

After rats completed seven daily MTP task training sessions and they made ≥ 60% correct responses on each lever, they were advanced to a version of the task wherein a delay was introduced between the sample and choice phases of each trial (DMTP). During each session, seven delay intervals were randomly intermixed across trials and each delay was presented for 16 trials (yielding a total of 112 trials/session). In the short delays phase of testing, delays ranged from 0 to 8 sec (0, 2, 3, 4, 5, 6, and 8 sec) and rats performed at least seven daily sessions before they were moved on to the last phase of DMTP. Those rats not performing two consecutive sessions with ≥ 75% correct averaged across delays remained on short delays for 1-2 additional sessions.

The long delay phase of DMTP used delays ranging from 0 to 24 sec (0, 2, 4, 8, 12, 18, and 24 sec) and daily sessions continued until rats met a performance criterion of ≥ 75% correct, collapsed across delays, for two consecutive sessions. They continued the long delay task for five “overtraining” sessions, or to a maximum of 14 total sessions. Lastly, rats were moved to a final testing phase that required delayed non-matching-to-position (DNMTP). Here, the rule was reversed such that during the choice phase, rats were required to select the lever not presented during the sample phase to earn a food pellet reinforcement. Long delays (0-24 sec) were utilized for the entirety of testing on the DNMTP task, which continued for 14 daily sessions.

### Data Analysis

Our data analysis plan was established a priori and is detailed in this study’s pre-registration (https://osf.io/tjxvd/). The effects of alcohol, THC, and drug combination exposure on body weight during the drug exposure period were determined by analyzing percent weight gain with two-way ANOVA (sex x treatment). Dose of alcohol (g/kg) were converted based on 10% ethanol consumed and body weight measured on each day. Intake of THC was confirmed for each rat by visual inspection that the presented cookie was completely consumed. In cases where rats did not consume an entire cookie, that dose was not included in the calculation of cumulative dose consumed for the entire drug exposure period. Group and sex differences in total intakes of SAC, EtOH, and THC were determined using two-way ANOVA (sex x treatment) and post hoc Tukey HSD tests were applied to reveal specific between-group differences following a significant main effect.

Performance on the DMTP and DNMTP tasks was assessed by calculating percent correct collapsed across delays for each session, calculating percent correct by delay, and by comparing group mean sessions to criterion. These analyses were chosen to investigate, respectively: potential differences in overall performance across the duration of testing on each task, differences in performance as the delay (and thus the difficulty) increased, and in the number of sessions it took animals in each group to meet pre-set criterion to progress to the next task. Performance during DMTP and DNMTP sessions were analyzed using a mixed ANOVA with drug treatment and sex as between-subjects factors, and session and/or delay as the within subject factor. Number of omissions was also analyzed separately for each delay and collapsed across delay.

Data from several animals were not included in the final analysis because they did not meet a priori inclusion criteria during the drug intake or behavioral assessment stages of the experiment. The total dose of THC given throughout the 18-day exposure period was 114 mg/kg. However, as the dose of THC increased to 3 and 5 mg/kg/cookie, some rats did not completely ingest one or both THC-laced cookies they were presented each day. Rats that missed > 29 mg/kg of THC were not included in data analysis, which led to exclusion of rates of 14 males and 5 females. During training and testing on the working memory task, 5 males and 4 females were excluded because they failed to pass the final training stage criteria for the MTP task, 1 male and 1 female were excluded from DNMTP analysis because of high rates of trial omissions (≥ 2 sessions with ≥ 15 omissions) after the rule change. In addition, 4 males had to be removed because they could not remain pair housed for the duration of the study due to the emergence of aggressive behavior. In total, 32 rats were removed from the final analysis.

## Results

### Fluid and drug consumption and bodyweight

After removing rats that did not meet the THC consumption criterion, there were no difference in the total dose consumed in the THC and COM groups for both sexes [group and sex effects: F(1, 47) = 0.36 and 1.57, p = 0.55 and 0.22, respectively; Fig 1A]. However, rats that consumed THC-laced cookies had significantly reduced intakes of 0.1% SAC and 10% ethanol solutions. As shown in Fig. 1B, female rats consumed significantly less SAC solutions [main effect of sex: F(1, 46) = 10.4, p < 0.003] and both sexes of THC consuming rats had significantly less total SAC intake than their respective controls [main effect of group: F(1, 46) = 11.2, p < 0.002; group x sex interaction: F(1, 46) = 0.08, p = 0.77]. As shown in Fig. 1C, there were no sex differences in ethanol intake [main effect of sex: F(1, 50) = 0.98, p = 0.33], but in both sexes co-use of THC significantly reduced EtOH intake [main effect of group: F(1, 3) = 40.53, p < 0.001]. THC consumption led to a reduced weight gain in male, THC-exposed animals, but co-use of ethanol alleviated this effect in the COM group [group effect: F(3, 96) = 4.12, p < 0.009 ; Fig 1D].

**Fig 1.**
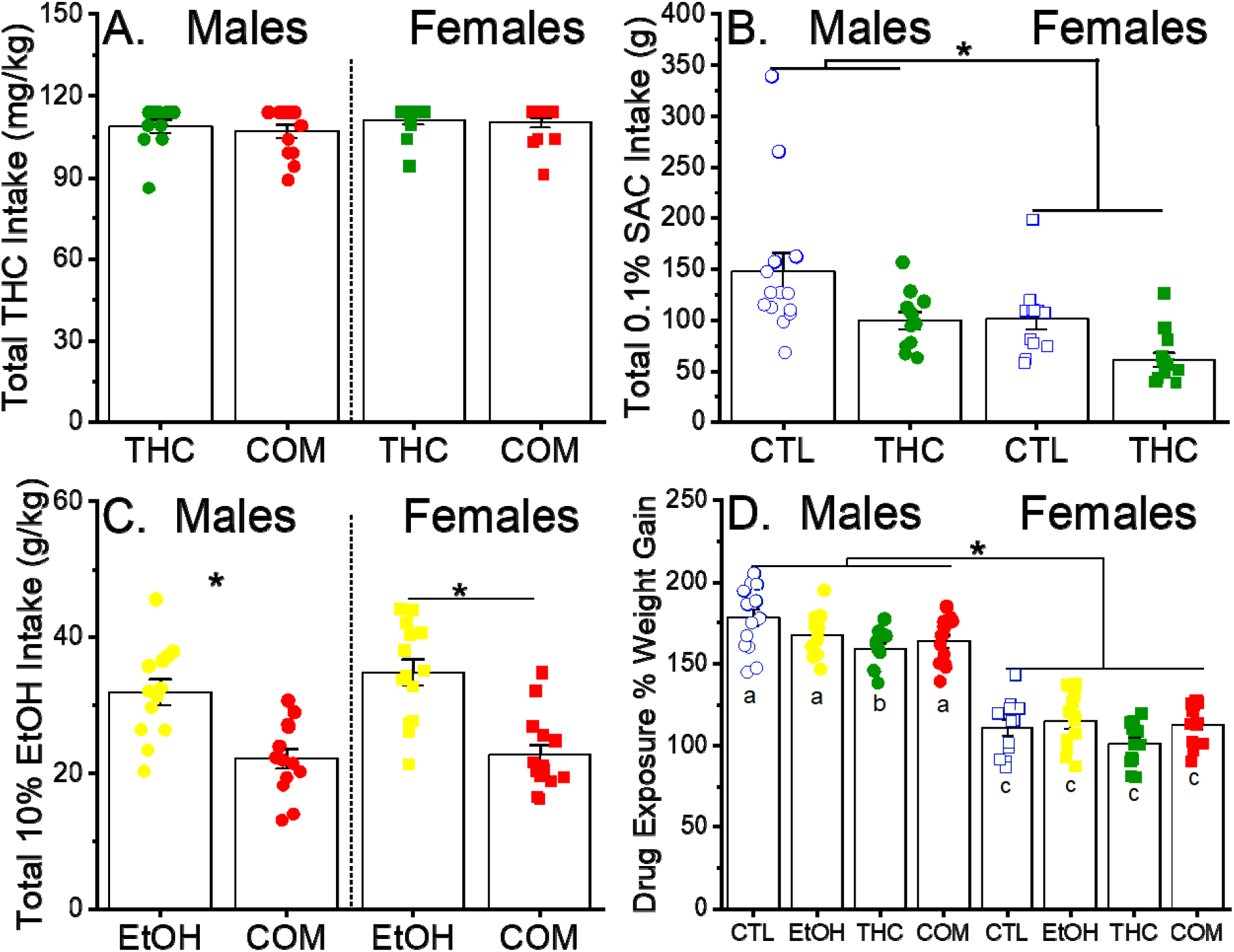
Drug intake and body weight gain during the adolescent treatment period. (A) Total intake of THC, which was offered on cookies at doses of 1,.5, 3.0 and 5.0 mg/kg/cookie, in rats that also drank 0.1% saccharin (SAC) or 0.1% SAC/10% ethanol (the THC and COM groups, respectively). The maximum possible intake of THC was 114 mg/kg. (B) Total intake of 0.1% SAC in rats that also ate vehicle-laced or THC-laced cookies (the CTL and THC groups, respectively). \**p* < 0.05, between treatment groups and sexes (C) Total intake of 0.1% SAC/10% ethanol in rats that also ate vehicle-laced or THC-laced cookies (the EtOH and COM groups, respectively). \**p* < 0.05, vs. EtOH within sex (D) Percent weight gain in rats from the beginning to the end of the treatment period (P28-P47). Different letters indicate a significant group difference based on post hoc tests, *p* < 0.05. Group size in males and females, respectively: CTL = 11 and 12, EtOH = 13 and 14, THC = 11 and 13, COM = 13 and 14.

### Plasma Levels of THC and Metabolites

Blood was collected from rats in the THC and COM groups 14-15 hours after their final self-administration session. Significant plasma levels of THC and THC-COOH were detected in rats of both sexes that were ingesting THC or both THC and ethanol (Table 2), but consistent with our previous work we did not detect significant levels of 11-OH-THC following oral consumption of THC (Nelson et al., 2019). In addition, there were no significant differences in THC levels between treatment groups, though data approached significance in females [males: F(1,22) = 0.08, p = 0.78; females: F(1,22) = 3.35, p = 0.08]. Female rats had significantly higher THC-COOH levels than their male counterparts, regardless of treatment [F(1,16) = 5.1, p = 0.038; Table 2].

**Table 2.** Plasma levels of THC and its major metabolites, 14-15 h after the last self-administration session in males and females allowed to consume THC and saccharin (THC) or THC and alcohol (COM).. Data are presented as mean (± SEM) concentrations (ng/mL) in each treatment group. \**p* < 0.05 versus males; *nd:* non-detectable

| Treatment | Sex | THC | 11-OH-THC | THC-COOH |
| --- | --- | --- | --- | --- |
| THC | Male | 1.53 (0.246) | *nd* | 0.536 (.170) |
|  | Female | 2.13 (0.52) | *nd* | 3.396 (1.062)* |
| COM | Male | 1.47 (0.31) | *nd* | 1.31 (0.602) |
|  | Female | 1.34 (0.207) | *nd* | 2.445 (0.600)* |

### Working memory task

Rats began testing on the short delay stage of the DMTP task at approximately P82 (35 days after the final drug exposure), and all animals completed testing by P120. Three-way ANOVA (sex x treatment x session or delay) revealed only modest differences in group performance at some stages of testing. At all stages of testing, there was a significant effect of the within-variable (session/delay), such that animals performed better in later sessions and on shorter delays, except as mentioned below in the first few sessions of DNMTP testing. When collapsed across delay, there was a main effect of sex [F(1,637) = 7.68, p = 0.002] on the DMTP short delay task, such that males performed significantly worse than females (Fig 2A). There were no significant effects of treatment, nor any significant interactions between sex, session, or treatment. When broken down by delay, there was a main effect of sex [F(1, 637) = 5.32, p = 0.021] observed at the early sessions of the DMTP short delay testing, with females outperforming their male counterparts (Fig 3A). Additionally, we observed a sex by treatment interaction [F(7, 637) = 2.76, p = 0.042], with post-hoc analyses confirming that female rats who consumed THC performed worse than their controls on the earlysessions of the short delay task (Fig 3). There were no significant main effects of treatment, or any other significant interaction effects. No significant differences in the mean days to meet criterion were found on the short delay DMTP task (Fig 2B). For the short delay stage, the mean number (± SEM) of omissions collapsed across groups was 2.61 ± 3.47, and there were no significant differences in omissions between any group in either the early or late sessions.

**Fig. 2.**
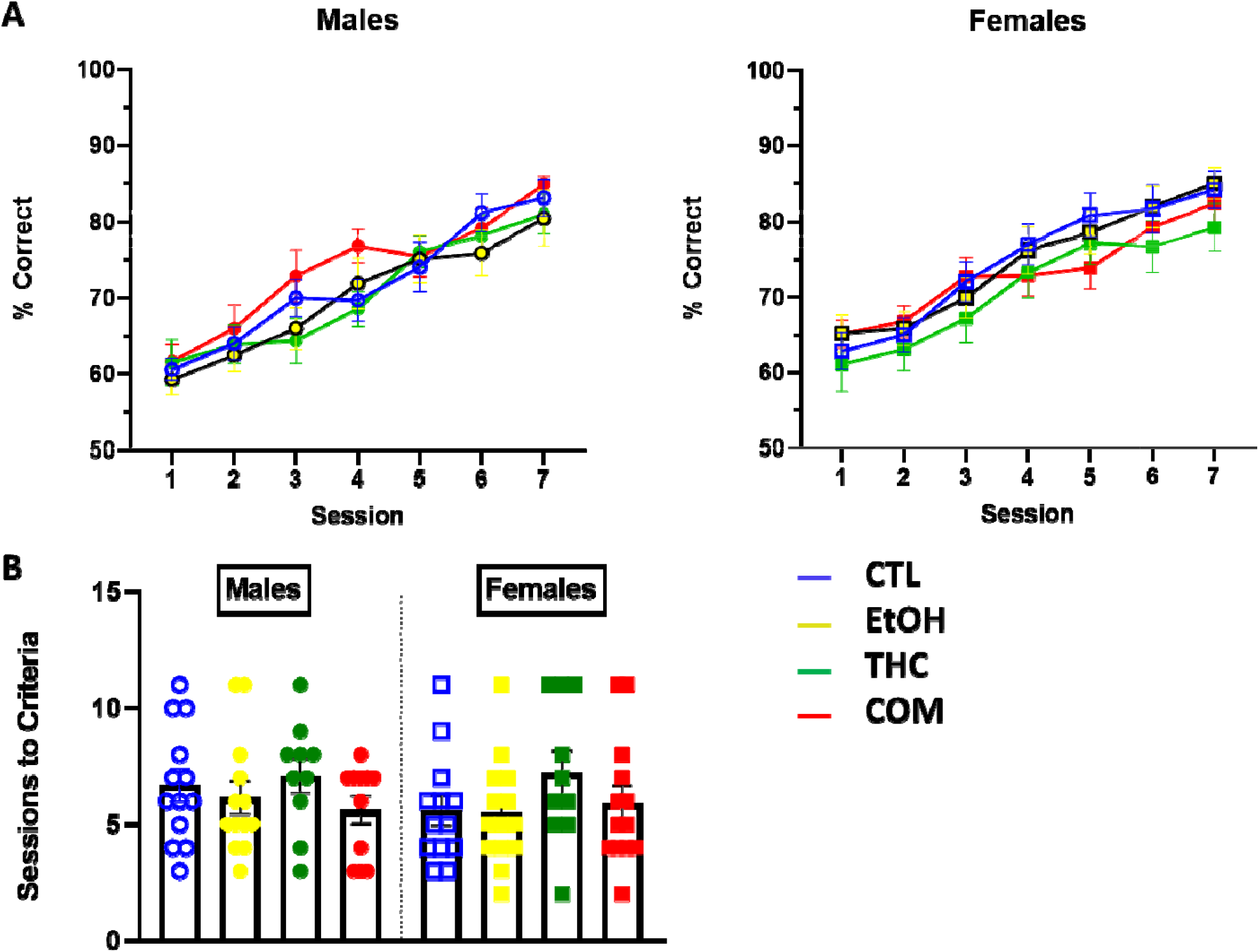
Performance on the short-delay (0-8s) portion of the delayed-matching-to-position task in control (CTL), ethanol (EtOH), THC, and combination (COM) animals (n= 9-14 per group). (A) Performance across the seven short-delay sessions that all rats completed, averaged across delay and presented separately for both sexes. (B). Average sessions to criterion (> 75% correct collapsed across delay for two consecutive days) for the short delay tasks.

**Fig. 3.**
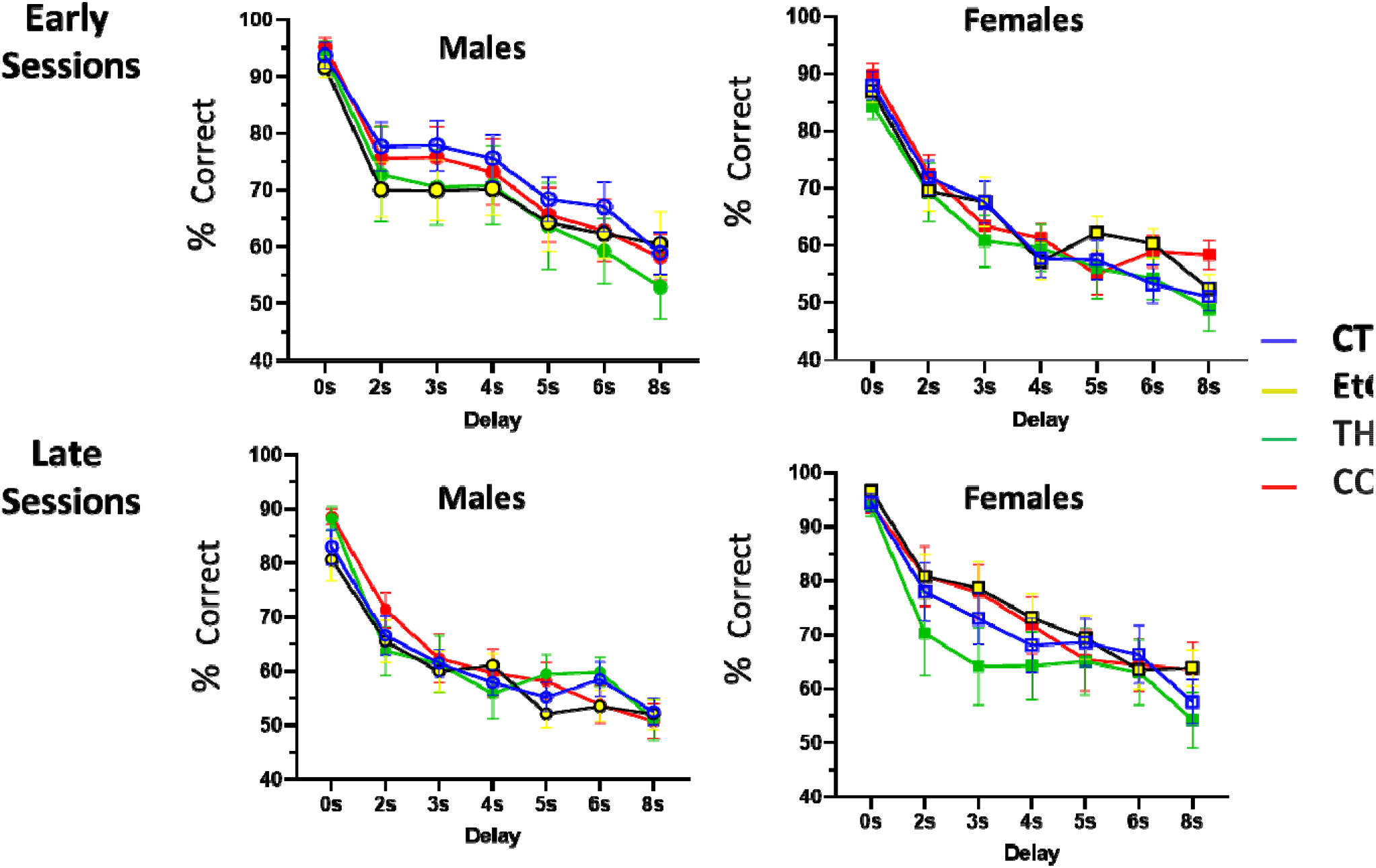
Performance during the beginning and end of the short-delay (0-8s) portion of the delayed-matching-to-position task in control (CTL), ethanol (EtOH), THC, and combination (COM) animals (n= 9-14 per group). Percent correct on early (average of sessions 1 and 2) and late (average of sessions 6 and 7) short-delay sessions broken down by delay and presented separately for both sexes.

On the long delay stage of the DMTP task, we found a sex-by-treatment interaction when data were collapsed across delay, such that males tended to perform better than their female counterparts in all treatment groups except the EtOH group, where females performed significantly better [F(3, 623) = 7.30, p < 0.001; Fig. 4A]. This sex-by-treatment effect persisted in early [F(3, 623) = 6.0, p < 0.001] and late [F(3,623) = 3.30, p = 0.019) sessions when broken down by delay (Fig 5), but no main effects were detected. There were no significant differences in the mean days to criterion on the long delay DMTP task (Fig 4B). For the long delay stage, the mean number (± SEM) of omissions collapsed across groups was 3.52 ± 4.11, and there were no significant differences in omissions between any group in either the early or late sessions.

**Fig. 4.**
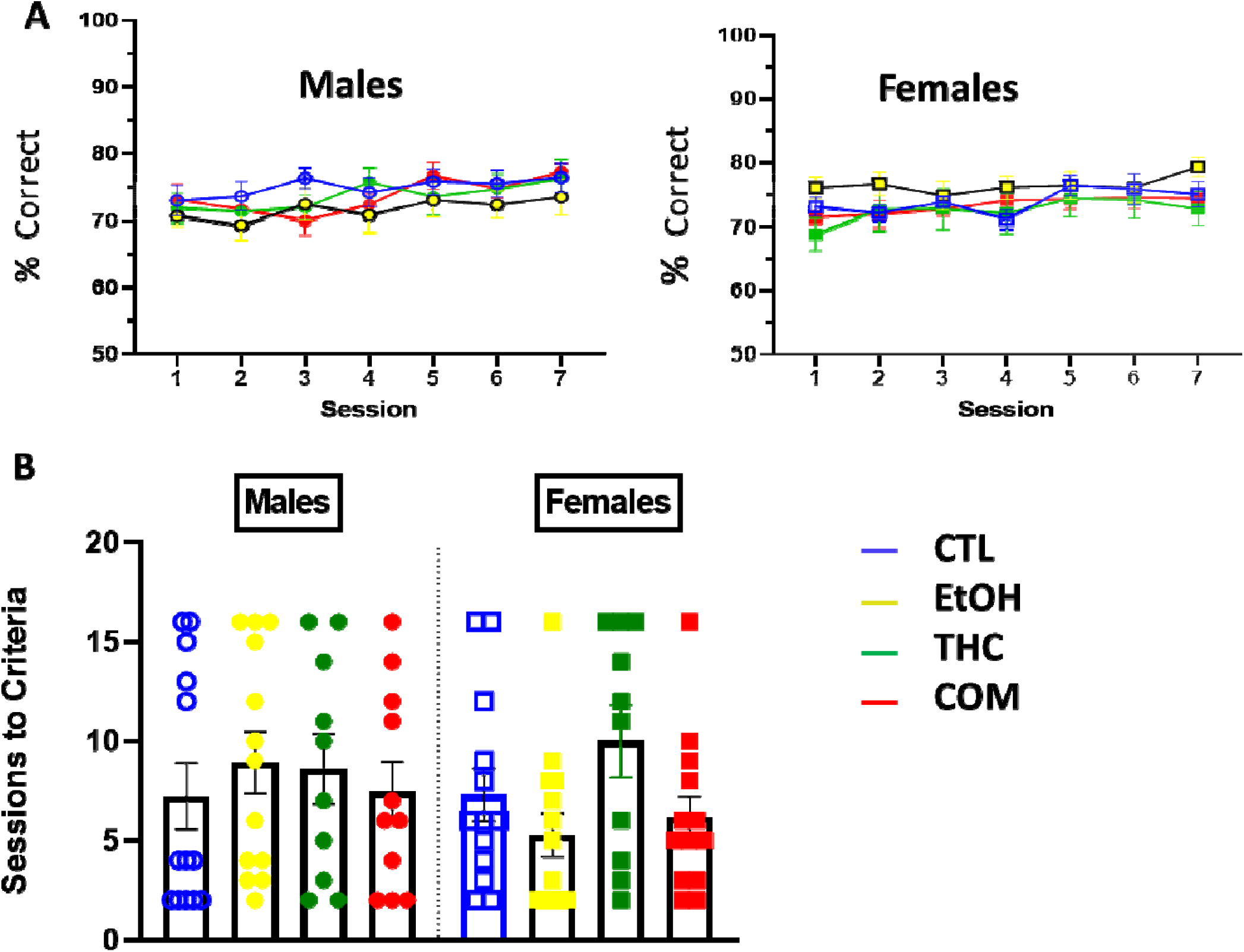
Performance on the long-delay (0-24s) portion of the delayed-matching-to-position task in control (CTL), ethanol (EtOH), THC, and combination (COM) animals (n= 9-14 per group). (A) Performance across the seven long-delay sessions that all rats completed, averaged across delay and presented separately for both sexes. (B) Mean number of sessions required for rats to meet criterion (>75% correct collapsed across delay for two consecutive days) for the long delay tasks.

**Fig. 5.**
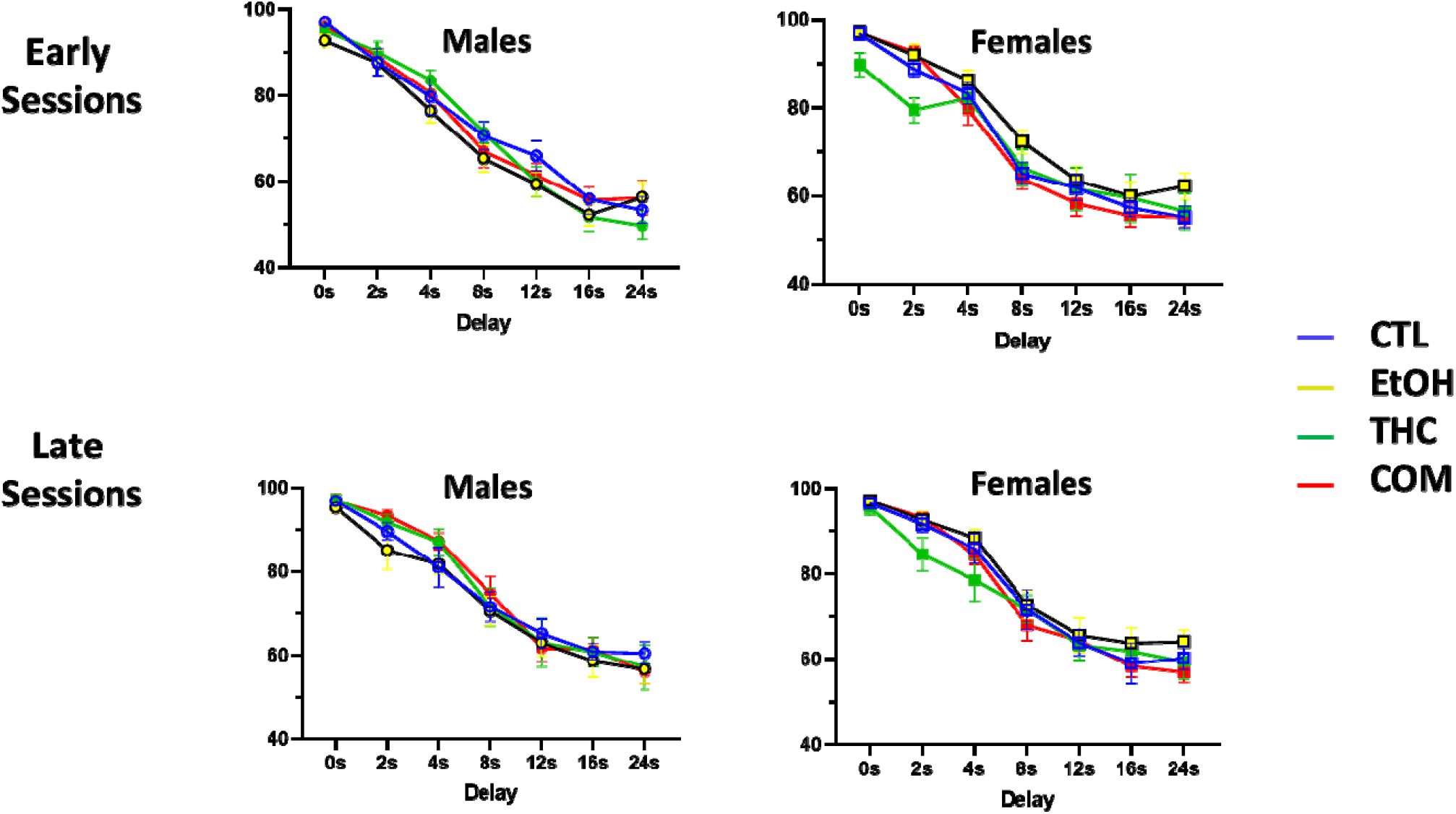
Performance during the beginning and end of the long-delay (0-24s) portion of the delayed-matching-to-position task in control (CTL), ethanol (EtOH), THC, and combination (COM) animals (n= 9-14 per group). Percent correct on early (average of sessions 1 and 2) and late (average of sessions 6 and 7) long-delay sessions broken down by delay and presented separately for both sexes.

For the DMTP tasks, accuracy decreased as delays increased (Fig 3 and Fig 5).When the rule was reversed to a non-match for the DNMTP task, performance decreased substantially in all groups and rats performed worse on the shorter delays than on longer delays in the early sessions (Fig. 6). Collapsed across delay, there was a significant effect of sex [F(1, 1274) = 5.30, p = 0.021] and a sex-by-treatment interaction [F(3, 1274) = 5.44, p = 0.001) such that females outperformed their male counterparts except in the THC group, where they performed worse (Fig 6A). The main effect of sex was not found when early and late sessions were broken down by delay, but the same sex-by-treatment interaction was significant in the late sessions [F(3,623) = 4.60, p = 0.003] (Fig 6B. The mean number (± SEM) of omissions collapsed across groups during the DNMTP task was 5.52 ± 6.11, and there were no significant differences between any group in either the early or late sessions. Because we did not find any significant group differences on the longer delays, we abandoned our a priori plan to analyze the potential role of proactive interference on performance deficits.

**Fig. 6.**
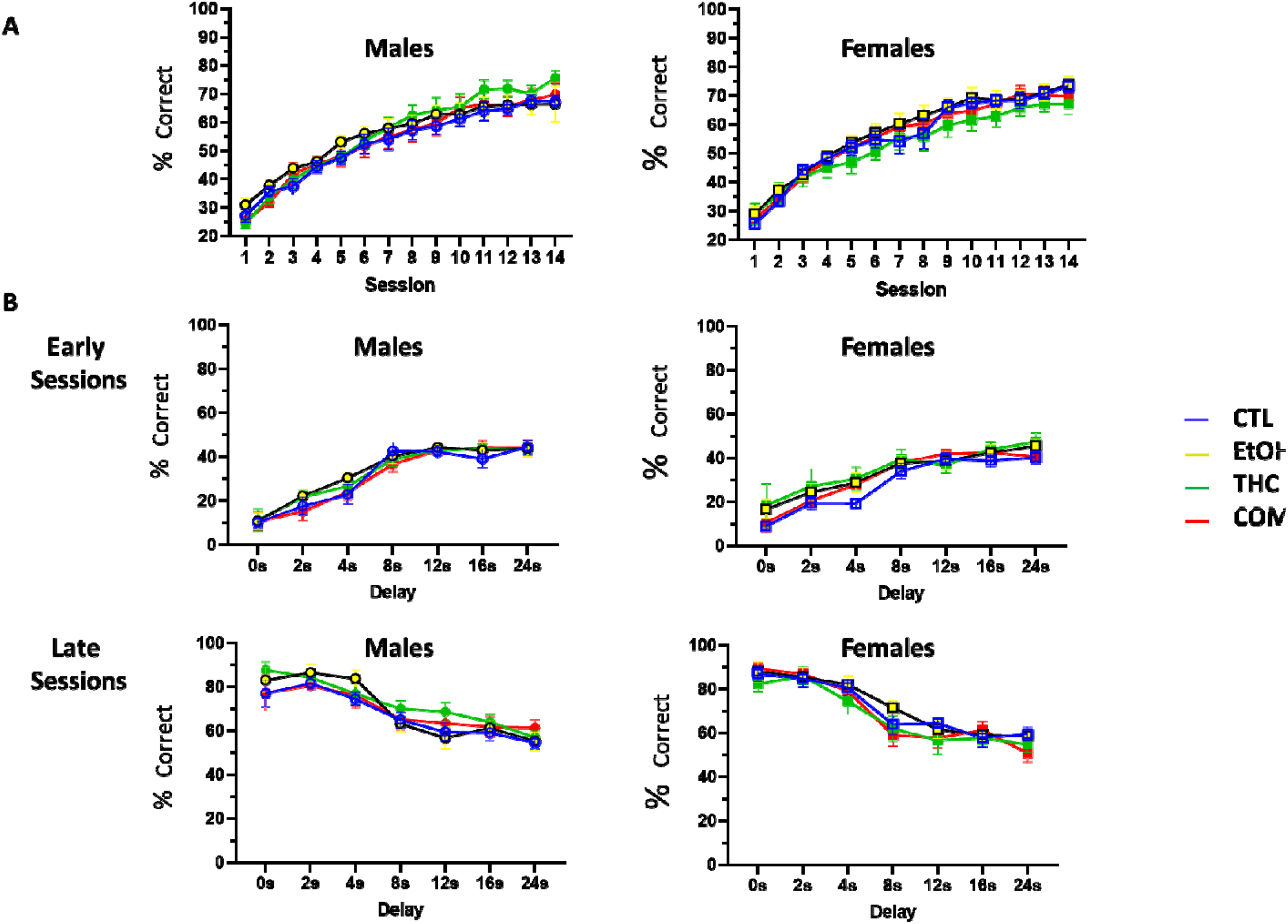
Performance on the delayed-non-matching-to-position task (0-24s), following the rule shift in control (CTL), ethanol (EtOH), THC, and combination (COM) animals (n= 9-14 per group). (A) Performance across the 14 sessions, averaged across delay and presented separately for both sexes. (B) Percent correct on early (average of sessions 1 and 2) and late (average of sessions 13 and 14) sessions broken down by delay and presented separately for both sexes.

## Discussion

Despite their simultaneous use being common, especially among adolescents and young adults (McCabe et al., 2021), the specific effects of adolescent co-use of THC and alcohol on long-term cognitive function has been studied infrequently. The available work in humans suggests that co-use of these drugs may lead to unique, more detrimental outcomes than the use of either drug alone (Noorbakhsh et al., 2020; Linden-Carmichael et al., 2020). To test whether this would be true in a rodent model, we allowed male and female adolescent rats to self-administer alcohol, THC, or both and tested their working memory in adulthood compared to control animals. As in our previous work, co-use of THC reduced voluntary consumption of ethanol. Analysis of blood levels of THC and its common metabolites following the final self-administration session revealed similar levels of THC in both sexes in EtOH and COM animals; however, female rats showed significantly higher THC-COOH levels than their male counterparts. In the DMTP task, we found a modest interaction effect of sex and treatment wherein females consistently outperformed their male counterparts except in the THC group. This effect was specific to the THC animals, and was not seen in our combination group animals, nor were there any consistent differences between any drug-treated groups and their same-sex controls. When the rule for reinforced response was reversed in the DNMTP task, animals in all groups took several days to adjust their responding. While we once again found an interaction effect of sex and treatment such that female THC-treated animals were the only group to be outperformed by their male counterparts, the lack of main effects of treatment suggest that neither THC, EtOH, or COM exposure significantly impacted behavioral flexibility in adulthood. Together, these data suggest that moderate, voluntary consumption of these drugs does not produce significant, long-lasting impairments in working memory or behavioral flexibility.

To model more closely the human phenomenon wherein most substance use is initiated during adolescence, we allowed our animals to voluntarily consume THC, ethanol, or both during the rat’s periadolescent period (P28-P47). While most pre-clinical studies of substance use utilize non-contingent exposure and thus force an animal to be exposed to a pre-determined amount of drug, studies have suggested that there are significant differences in behavior and drug-induced neural adaptations in animals allowed to willingly consume a given drug (Jacobs et al., 2003; Schweppe et al., 2020). This includes changes to their willingness to self-administer other drugs, e.g. how much alcohol a rat will drink after willingly consuming THC (Nelson et al., 2019a). Consistent with our previous work (Nelson et al., 2019a; Nelson et al., 2019b), THC consumption reduced weight gain in male rats, but co-use of alcohol attenuated this effect. While some human literature suggests that simultaneous use of THC leads to greater alcohol intake (Linden-Carmichael et al., 2017; Linden-Carmichael et al., 2020), we found that rats consuming THC reduced their saccharin or ethanol intake in the THC and combined groups, respectively, which is consistent with our previous findings (Nelson et al., 2019a). In another study where male adolescent rats were allowed to consume alcohol either in isolation or on days where they were also exposed to vaporized THC, Hamidullah et al. (2021) found that rats increased their alcohol intake on vehicle exposure days and decreased alcohol intake on THC-exposure days. This is consistent with findings in cannabis-dependent humans during cannabis abstinence (Allsop et al., 2014; Lucas et al., 2016), as well as in the National College Health Assessment Survey-II which has shown a decrease in reported binge alcohol consumption in states where recreational cannabis use has been legalized (Alley et al., 2020). Together, these findings show that THC may have an acute suppressive effect on alcohol drinking, suggesting a “replacement effect” wherein more alcohol is consumed to make up for the lack of THC effects. The long-term effects of THC use on alcohol consumption are not well understood and need to be studied further, particularly in those who initiate co-use during adolescence.

Both alcohol and THC use alone in adolescence has been demonstrated to impair cognition (Spear, 2018; Bara et al., 2021). The duration and severity of cognitive impairments, as well as if any impairments are found at all, are influenced by numerous factors including sex, age, route of administration, dose and pattern of drug exposure, withdrawal time prior to testing, and the cognitive task assessed. For example, Cha et al. (2007) tested rats in a Morris water maze 30 min after an i.p. injection of 5 mg/kg THC and reported evidence of impaired spatial learning in adolescents and females compared to adults and males, respectively. However, once daily exposure to this dose of THC for 30 days had no effect on task performance when assessed 28 days after the final drug injection. A study that utilized an escalating dose, non-contingent exposure paradigm, reported that THC slowed, but did not prevent, acquisition of an associative learning task when training occurred two weeks after the final drug exposure. Once animals had learned the task, there was no effect of THC on performance (Abela et al., 2019). Similarly, the effects of alcohol on cognitive function are highly variable and dependent on the methodology utilized. Rats exposed repeatedly to vaporized ethanol during adolescence show deficits in behavioral flexibility when they are tested in adulthood (Gass et al., 2014), but adult male rats who self-administer a low to moderate amount of alcohol exhibit improvement in behavioral flexibility (Cacace et al., 2011). In the current study, male and female rats were allowed to self-administer alcohol, THC, both, or their vehicles over a periadolescent timeframe and then tested on a working memory and behavioral flexibility task in adulthood. The total drug consumed corresponded to low- or moderate of intake, with several rats removed from the study due to refusal to ingest the higher doses of THC towards the end of the treatment period. We found only modest effects of THC treatment that was specific to female rats, and most consistently this was only in comparison to their male counterparts that also consumed THC. Notably, male and female rats undergo pubertal onset at different time points, with males maturing more slowly than females. Differences in maturational stage versus differences in chronological age during exposure to drugs like THC and alcohol remains an issue whenever animals of both sexes are tested. Further analysis would be needed to investigate whether or not these differences in maturational state during exposure may play a role in sex-specific effects of one or both of these substances.

Interest in understanding the risks conferred by patterns of simultaneous use has been increasing (Lee et al., 2022) alongside the greater incidence of co-use in young adults (McCabe et al., 2021). Most of the research in the human literature focuses on acute effects of simultaneous use, and specifically on the immediate consequences that are self-reported by users of these drugs (Lee et al., 2022). When longitudinal studies are conducted, they most frequently compare “heavy” users of one or both drugs to non-using controls, and even then the criteria for heavy frequency of use is inconsistent, making it difficult to paint a comprehensive picture (Karoly et al., 2020). With less heavy, “recreational” use of these drugs, there is some evidence in humans that the effects of alcohol can be attenuated by cannabis use (Mahmood et al., 2010; Infante et al., 2018), similar to findings in rodents (Hamidullah et al., 2021). Here, we aimed to model low to moderate intake of these drugs that is more consistent with non-disordered use of these substances since this is the type of use that is more common in society. According to estimates from the 2021 National Survey on Drug Use and Health, about 15% of Americans between the ages of 12-20 are current users of alcohol and only about 2% can be classified as “heavy” users (SAMHSA, 2021). Similar data for adolescent THC use is less concisely reported, perhaps given a lack of an accepted cut-off for “heavy” use of marijuana; however, a study in U.S. high school students reported that about 22% were current users of cannabis, with most of them (about 59%) using the drug 1-9 times a month (Jones et al., 2020). Thus, a majority of of the drug-using adolescent population are not taking heavy amounts of either drug and this emphasizes the need to understand the potential impact of more moderate use on the developing brain. In the current study, we did not observe a significant impact of alcohol or THC use, alone or in combination, but more studies that utilize moderate and voluntary intake in laboratory animals is necessary to increase our understanding of what, if any, unique impact results from co-use. A comprehensive understanding of the potential impacts of co-use of alcohol and THC, especially during adolescence, is important for advising social policies as well as treatment strategies if drug use becomes disordered.

## Acknowledgements

The authors thank David Tkac and Julia Minkevitch for their technical assistance with these experiments. This work was supported in part by funding from The National Institute on Drug Abuse (R21 DA 045175) awarded to NCL and JMG.

